# Effectiveness of fixation methods for wholemount immunohistochemistry across cellular compartments in chick embryos

**DOI:** 10.1101/2024.03.23.586361

**Authors:** Camilo V. Echeverria, Tess A. Leathers, Crystal D. Rogers

**Author notes:** These authors contributed to this manuscript equally (Camilo V. Echeverria, Tess A. Leathers).

## Abstract

The choice of fixation method significantly impacts tissue morphology and protein visualization after immunohistochemistry (IHC). In this study, we compared the effects of paraformaldehyde (PFA) and trichloroacetic acid (TCA) fixation prior to IHC on chicken embryos. Our findings underscore the importance of validating fixation methods for accurate interpretation of IHC results, with implications for antibody validation and tissue-specific protein localization studies. We found that TCA fixation resulted in larger and more circular nuclei compared to PFA fixation. Additionally, TCA fixation altered the appearance of subcellular localization and fluorescence intensity of various proteins, including transcription factors and cytoskeletal proteins. Notably, TCA fixation revealed protein localization domains that may be inaccessible with PFA fixation. These results highlight the need for optimization of fixation protocols depending on the target epitope and model system, emphasizing the importance of methodological considerations in biological analyses.

## 1. INTRODUCTION

Immunohistochemistry (IHC) stands as a cornerstone to visualize cell and tissue-level phenomena, revealing molecular interactions and protein localization within biological specimens. Central to this technique is the process of fixation, crucial for preserving tissue morphology and antigenicity. Here, we explore the comparative efficacy of two prevalent fixatives—paraformaldehyde (PFA) and trichloroacetic acid (TCA)—prior to immunohistochemical analysis. The study delves into the respective impacts of each fixative method and time length on antigen localization, cellular morphology, and signal intensity. Our goal is to unravel the nuanced effects on the quality and reliability of IHC outcomes and potential differences between the two methods. IHC can lead to variable results depending on the tissue sample used, antibody efficacy, and antigen type and localization (Ayoubi et al., 2023). Prior studies identified that specific fixation methods are necessary to visualize proteins that are localized to different sub-cellular regions or cellular structures (Feng et al., 2021; Hua and Ferland, 2017). Through a systematic investigation, we provide comprehensive insights into how the choice of fixative can alter results, which may empower researchers in optimizing immunohistochemical protocols for enhanced accuracy and reproducibility in biological analyses. Specifically, here we analyze the outcomes of fixing wholemount *Gallus gallus* (chicken) embryos with PFA and TCA without antigen retrieval and use IHC to identify how those methods alter the localization and fluorescence intensity of proteins that are normally found in the nucleus, cytoplasm, and cell membrane.

In developmental biology, investigating protein localization changes in vertebrate embryos using IHC offers a profound understanding of intricate molecular processes governing embryogenesis. At minimum, IHC can provide basic details of cell and tissue types in which a protein is expressed, but the technique can also offer insight into dynamic cellular and subcellular localization changes of specific proteins across developmental stages. The selection of fixation methods significantly influences the accuracy and fidelity of developmental studies. Given the delicate nature of embryonic tissues, a multitude of IHC studies using embryonic tissues use aldehyde fixation in the form of formaldehyde, formalin or PFA (Table 1). PFA is often favored for embryonic specimens due to its ability to cross-link proteins and amines in DNA and RNA, thus preserving tissue architecture and maintaining structural epitopes (Stumptner et al., 2019). Upon contact with tissue, PFA undergoes hydrolysis to form formaldehyde, its active component, and this reactive aldehyde efficiently crosslinks proteins via amino acid bridges (Klockenbusch and Kast, 2010; Nadeau and Carlson, 2007; Solomon and Varshavsky, 1985). The ability of PFA to create stable crosslinks makes it the fixative agent of choice to preserve structural epitopes for subsequent microscopic analysis and downstream experimentation.

**Table 1.** Common fixative methods used for developing vertebrate embryos.

| Fixative Chemical | Chicken | Zebrafish | Frogs |
| --- | --- | --- | --- |
| **Aldehyde-based** | (Balint and Csillag, 2007; Carro et al., 2013; Chacon and Rogers, 2019; Rogers et al., 2013a; Rogers et al., 2013b) | (Hammond-Weinberger and ZeRuth, 2020; Macdonald, 1999; Shimomura et al., 2007) | (Acton et al., 2005; Fagotto and Brown, 2008; Kurth et al., 2010; Lee et al., 2008; Robinson and Guille, 1999; Tao et al., 2007) |
| **Alcohol-based** |  |  | (Lee et al., 2008; Ossipova et al., 2022; Ossipova and Sokol, 2021) |
| **Acid-based** |  | (Lasseigne et al., 2021; Macdonald, 1999; Martin et al., 2022) |  |
| **Other** |  |  | (Acton et al., 2005) |

Conversely, TCA fixation, known for its permeabilization and dehydration, presents an alternative with potential benefits to access hidden epitopes in embryos but is used less frequently (Lee et al., 2008; Martin et al., 2022). Upon application, TCA penetrates tissues and promptly precipitates proteins by causing their denaturation and aggregation through acid-induced coagulation, which may enhance or deter the ability of antibodies to bind to specific antigens depending on their target (Rajalingam et al., 2009). The acidic nature of TCA and high precipitation capacity result in rapid and robust fixation, preserving tissue architecture by solidifying cellular constituents and preventing enzymatic degradation (Hao et al., 2015; Lakatos and Jobst, 1992). While TCA fixation may alter some protein structures due to its denaturing effects, this can be beneficial when used against bulky or hidden epitopes in subsequent histochemical and immunohistochemical analyses.

Visualizing various types of proteins within cells demands a careful approach, considering the diverse subcellular localizations and unique tertiary structures they may possess. For proteins localized to distinct subcellular regions such as the nucleus, cytoplasm, or plasma membrane, fixation methods must cater to the preservation of these specific environments. Fixatives like PFA are adept at maintaining the intricate membranous structures and spatial organization within the cytoplasm or plasma membrane. In addition, for proteins residing in the nucleus, fixation methods that effectively permeate nuclear membranes and preserve nuclear morphology become imperative. Moreover, proteins with intricate tertiary structures, such as those forming multimeric complexes or undergoing post-translational modifications, often necessitate fixation techniques that maintain these delicate interactions. Thus, tailoring fixation methods according to subcellular localization and protein tertiary structure becomes pivotal in accurately visualizing diverse protein populations within cells.

The selection of fixation methods in IHC poses a delicate balance between tissue preservation and antibody penetration. While certain fixatives excel in preserving tissue architecture and antigenicity, their robustness might hinder the penetration of certain antibodies into the tissue, limiting the accessibility to targeted antigens. Conversely, fixation methods optimized for better antibody penetration might compromise tissue integrity and antigen preservation, which can alter the ability to use these tissues for downstream processing. Achieving an optimal equilibrium between these two facets is crucial to ensure comprehensive visualization of antigens within tissues, balancing the preservation of structural integrity with the facilitation of antibody access for accurate and reliable analyses.

With this study, we demonstrate the efficacy of PFA and TCA fixation methods specifically in the context of visualizing avian embryonic development using IHC, shedding light on their distinct impacts on tissue preservation and antigen detection to aid researchers in selecting the most suitable approach for developmental investigations. Here, we identify that TCA fixation methods may be optimal to visualize cytosolic microtubule subunits and membrane-bound cadherin proteins, but that TCA is subpar to visualize nuclear-localized transcription factors after IHC. In contrast PFA provides adequate signal strength for proteins localized to all three cellular regions but is optimal for maximal signal strength of nuclear-localized proteins.

## 2. MATERIALS AND METHODS

### Collection and staging of chicken embryos

Fertilized chicken eggs were obtained from UC Davis Hopkins Avian Facility and incubated at 37°C to the desired stages according to the Hamburger and Hamilton (HH) staging guide. After incubation, embryos were dissected out of eggs onto Wattman filter paper and placed into room temperature Ringer’s Solution. Embryos were then fixed using one of the methods listed below prior to IHC.

### Fixation Methods

#### Paraformaldehyde

Paraformaldehyde (PFA) was dissolved in 0.2M phosphate buffer to make 4% (w/v) stock solution and was stored at -20°C prior to use and was thawed fresh before use. Embryos were fixed at room temperature with 4% Paraformaldehyde (PFA) for 20 minutes (20 m). After fixation, embryos were washed in 1X Tris-Buffered Saline (TBS; 1M Tris-HCl, pH 7.4, 5M NaCl, and CaCl_2_) containing 0.5% Triton X-100 (TBST+Ca^2+^) or 1X Phosphate Buffered Saline (PBS) containing 0.1-0.5% Triton X-100 (PBST).

#### Trichloroacetic Acid

Trichloroacetic acid (TCA) was dissolved in PBS to make 20% (w/v) stock solution and stored at -20°C prior to use. It was then thawed and diluted to 2% concentration with PBS fresh before use. Embryos were fixed at room temperature with 2% Trichloroacetic Acid (TCA) in 1X PBS for 1-3 hours (hr). After fixation, embryos were washed in TBST+Ca^2+^ or PBST.

### Immunohistochemistry

After fixation, embryos were washed with the same buffer in which the fixative was diluted (see above) and wholemount IHC was performed. To block against non-specific antibody binding, embryos were incubated in PBST or TBST+Ca^2+^ containing 10% donkey serum (blocking solution) for 1 hour at room temperature or overnight (12-24 hr) at 4°C. Primary antibodies were diluted in blocking solution at indicated dilutions (Table 2) and embryos were incubated in primary antibodies for 72–96 hours at 4ºC. Multiple antibodies from the study have previously been validated in cell lines or embryos. After incubation with primary antibodies, whole embryos were washed in PBST or TBST+ Ca^2+^, then incubated with AlexaFluor secondary antibodies diluted in blocking solution (1:500) overnight (12-24 hr) at 4ºC. They were then washed in PBST or TBST+ Ca^2+^ as the final step before imaging for TCA-fixed embryos or washed with TBST+ Ca^2+^ post-secondary incubation and were post-fixed with PFA for 1 hour at room temperature for PFA-fixed embryos.

**Table 2.** Antibodies used in study.

| Antibody target | Company and catalog # | Dilution | Isotype | Protein type/Expected localization | Immunogen |
| --- | --- | --- | --- | --- | --- |
| PAX7 | DSHB (PAX7) | 1:5-1:10 | Mouse IgG1 | TF/Nucleus | Amino acids 418-427 |
| SNAI2 | Cell signaling, mAb9585 | 1:200 | Rabbit IgG | TF/Nucleus | Recombinant human Slug protein |
| SOX9 | Millipore Sigma, AB5535 | 1:500 | Rabbit IgG | TF/Nucleus | Epitope: C-terminal KLH-conjugated linear peptide corresponding to the C-terminal sequence of human Sox9 |
| ECAD | BD<br>Transduction<br>Laboratories<br>61081 | 1:500 | Mouse IgG2a | Cell<br>adhesion/Membrane | Amino acids 735-883 |
| NCAD | DSHB (NCAD),<br>MNCD2 | 1:5 | Rat IgG1 | Cell<br>adhesion/Membrane | Chicken N-cadherin |
| TUBA4A | Sigma, T6199 | 1:250 | Mouse IgG1 | Cytoskeleton/Cytosol | C-terminal end of the α-tubulin<br>isoform (amino acids 426-430) |
| TUBB2A | Abnova,<br>H00007280-M03 | 1:250 | Mouse IgG2a | Cytoskeleton/Cytosol | TUBB2A (full-<br>length recombinant protein with<br>GST tag |
| TUBB3 | R&D Systems,<br>MAB1195 | 1:250 | Mouse IgG2a | Cytoskeleton/Cytosol | Rat brain-derived microtubules |

### Cryosectioning

Following whole embryo imaging, embryos were prepared for cryosectioning by incubation with 5% sucrose in PBS (30 mins to 1 hr at room temperature), followed by 15% sucrose in PBS (3 hr at room temperature to overnight at 4ºC), and then in 10% gelatin with sucrose in PBS for 3 hr to overnight at 38-42ºC. Embryos were then flash frozen in liquid nitrogen and were sectioned in an HM 525 NX Cryostats, Epredia, Richard-Allan Scientific in 16 µm sections.

### Microscopy

Fluorescence images were taken using Zeiss ImagerM2 with Apotome.2 and Zen software (Karl Zeiss). Whole embryos were imaged at 10X (Plan-NEOFLUAR 10X/0,3 420340-9901) and transverse sections were imaged at 20X (Plan-APOCHROMAT 20X/0,8 420650-9901) with Apotome optical sectioning. Images were adjusted for brightness and contrast uniformly across the entire image in Adobe Photoshop in accordance with journal standards.

### Nuclear Pixel Intensity

To find the pixel intensity from one side of the nuclear membrane to the other, 10 nuclei from 5 different embryos per marker were analyzed using NIH ImageJ/Fiji with the Dynamic ROI Profiler plugin. These measurements were performed on images converted to grayscale with manual brightness and contrast adjustments through Photoshop. Using ggplot in R studio, the averaged pixel intensity across the length of the nuclei is plotted with a 95% confidence interval.

### Fluorescence intensity analysis

Fluorescence was quantified using NIH ImageJ/Fiji by averaging the relative intensity of tissue-specific regions in section images of chicken embryos. Sections were converted to grayscale, and contrast was adjusted automatically for each section using Adobe Photoshop. The grayscale images were analyzed using the rectangle tool to quantify the differences in fluorescence between neural crest (NC) cells and cells within the neural tube (NT). The rectangle tool was used to select four 0.30 x 0.30-pixel-size regions where NC cells are present and within the NT. The measurements obtained through Fiji included the “area,” “area of integrated intensity,” and “mean grey value.” The background was also measured for fluorescence. The corrected total cell fluorescence (CTCF) was calculated by subtracting the “area integrated density” from the product of the “area” of a selected region of interest and the “mean gray value” of the background, averaged out the values obtained from each region, then graphed. This was repeated when comparing the non-neural epithelium (NNE) and the cranial mesenchyme.

## 3. RESULTS

### TCA fixation increases nuclear area, circularity, and alters protein localization compared to PFA

To identify if fixation methods affected the general tissue structure, we tested the various fixation methods (4% PFA for 20 min, 2% TCA for 1 hr and 3 hr) in Hamburger Hamilton stage 9 (HH9) chicken embryos. Embryos were collected as described in the methods and fixed in their respective fixatives for 20 minutes, 1 hr, or 3 hrs. After fixation, IHC was performed using the antibodies in Table 2 and embryos were stained using the nuclear DNA stain, 6-diamidino-2-phenylindole (DAPI). Although no overt differences were observed in wholemount images of HH8 embryos (Figure 1A-C), after cryosectioning, TCA-fixed neural tubes (NT) maintained their morphological structure better, and all nuclei were significantly larger than nuclei from PFA-fixed embryos (Figure 1D-F). To quantify the differences in cell area or circularity, nuclei from 2-4 regions in the NT and collectively migrating neural crest (NC) cell regions were outlined using FIJI and assessed for both area and circularity. Using the Mann-Whitney U test to compare the anatomical differences between the TCA and PFA-fixed samples identified that both NT and NC nuclei had significantly larger areas and were more circular after TCA fixation (Figure 1G-J). Compared to fixation using 4% PFA, 2% TCA fixation resulted in NT nuclei with a larger area in embryos HH9 and older (147% larger (1 hr) and 137% larger (3 hr), P ≤ 0.0001). In collectively migrating NC cells, the average area of the nuclei was increased (148% larger (1 hr) and 140% larger (3 hr), P ≤ 0.0001). Using Fiji, circularity was measured in NT and NC cells. The formula for circularity is 4pi(area/perimeter^2). A value of 1.0 indicates a perfect circle. In NT cells, the PFA-fixed nuclei had an average circularity score of 0.69, while the TCA-fixed cells had scores of 0.83. In NC cells, the PFA-fixed nuclei had an average circularity score of 0.81, while the TCA-fixed cells had scores of 0.85 and 0.86. These morphological changes supported our observation that nuclei appeared more diffuse in 2% TCA-fixed samples compared to 4% PFA fixation (Figure 1 and 2).

**Figure 1.**
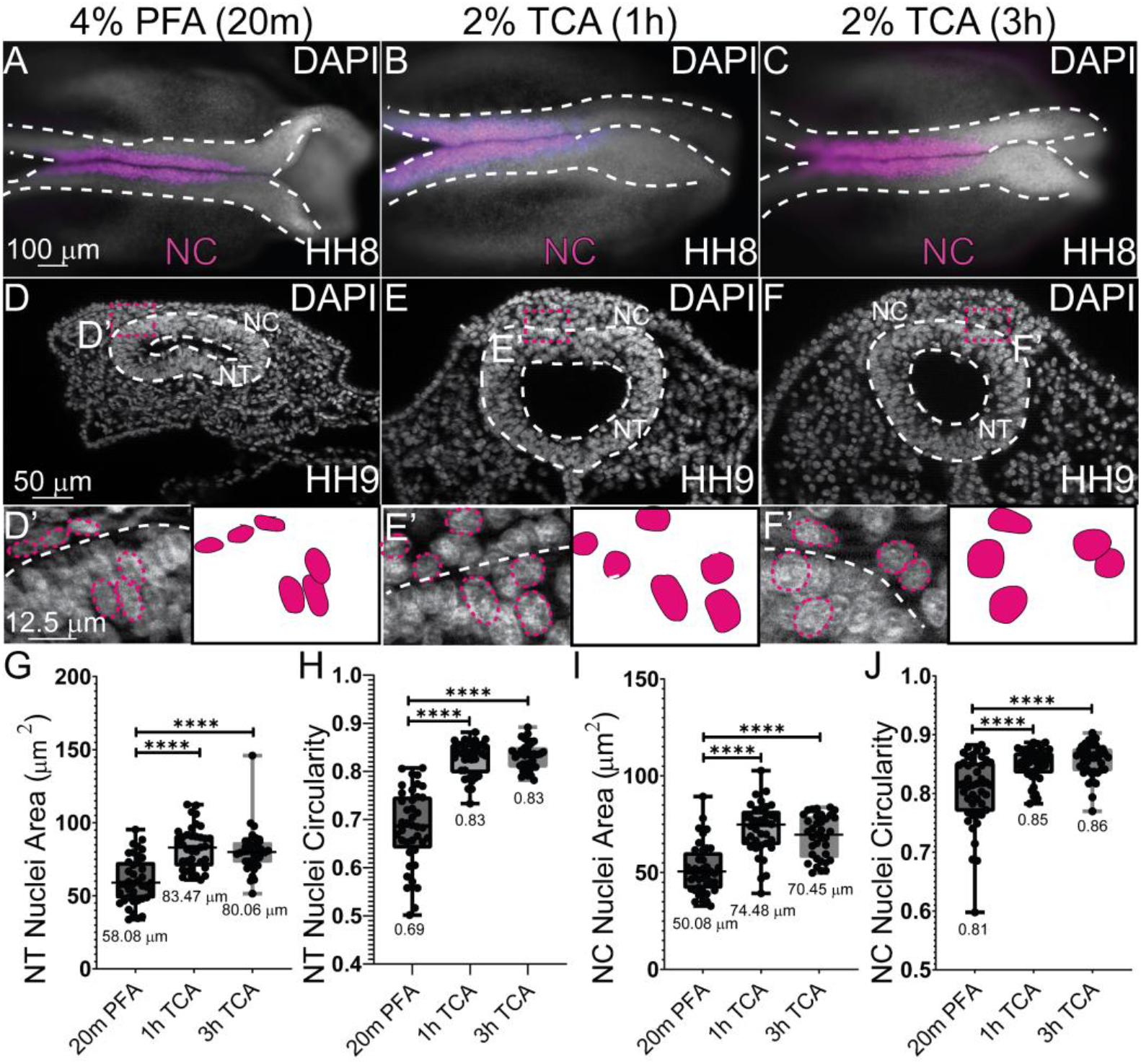
TCA fixation alters nuclear area and circularity. (A-C) Wholemount chicken embryos staged HH8-9 after IHC for NC marker (SOX9) and DAPI staining (cranial region, anterior to the right). Embryos were fixed in (A, D, D’) 4% PFA for 20m (B, E, E’), 2% TCA for 1h (C, F, F’), and 2% TCA for 3h. (D-F) Transverse cryosections from HH9 embryos. (D’-F’) Zoomed in regions from (D-F) with NT outlined in white dashed lines and select nuclei in pink dashed lines. (G) The area of the NT nuclei are significantly different between PFA and the two TCA treatment incubations. The mean NT nuclei area for the PFA treated is 58.08 μm^2^ (*n*=40), 1h TCA is 83.47 μm^2^ (*n*=36), and 3h TCA is 80.06 μm^2^ (*n*=36), P ≤ 0.0001. (H) NT nuclei circularity was measured, and significant differences were observed between PFA with a mean of 0.89 (*n*=40) and 1h TCA with a mean of 0.83 (*n*=36) and 3h TCA treatment with a mean of 0.83 (*n*=36), P ≤ 0.0001. (I) NC cell nuclei area was measured and PFA nuclei had significantly less area, with a mean of 50.08 μm^2^(*n*=40), 1h TCA with 74.48 μm^2^ (*n*=36), and 3h TCA with a mean of 70.45 μm^2^ (*n*=36) P ≤ 0.0001. (J) NC cell nuclei circularity differed PFA and the 1hr and 3hr TCA fixation. The mean circularity for PFA NC nuclei circularity was at 0.81, the mean for 1hr TCA was 0.85, and the mean for 3h TCA is 0.86, *n*=36, P ≤ 0.0001. One-way ANOVA with the Mann-Whitney test was used to determine the significance. The scale bar for the wholemount images is 100 μm, the transverse sections is 50 μm, and the high magnification transverse section is 12.5 μm.

**Figure 2.**
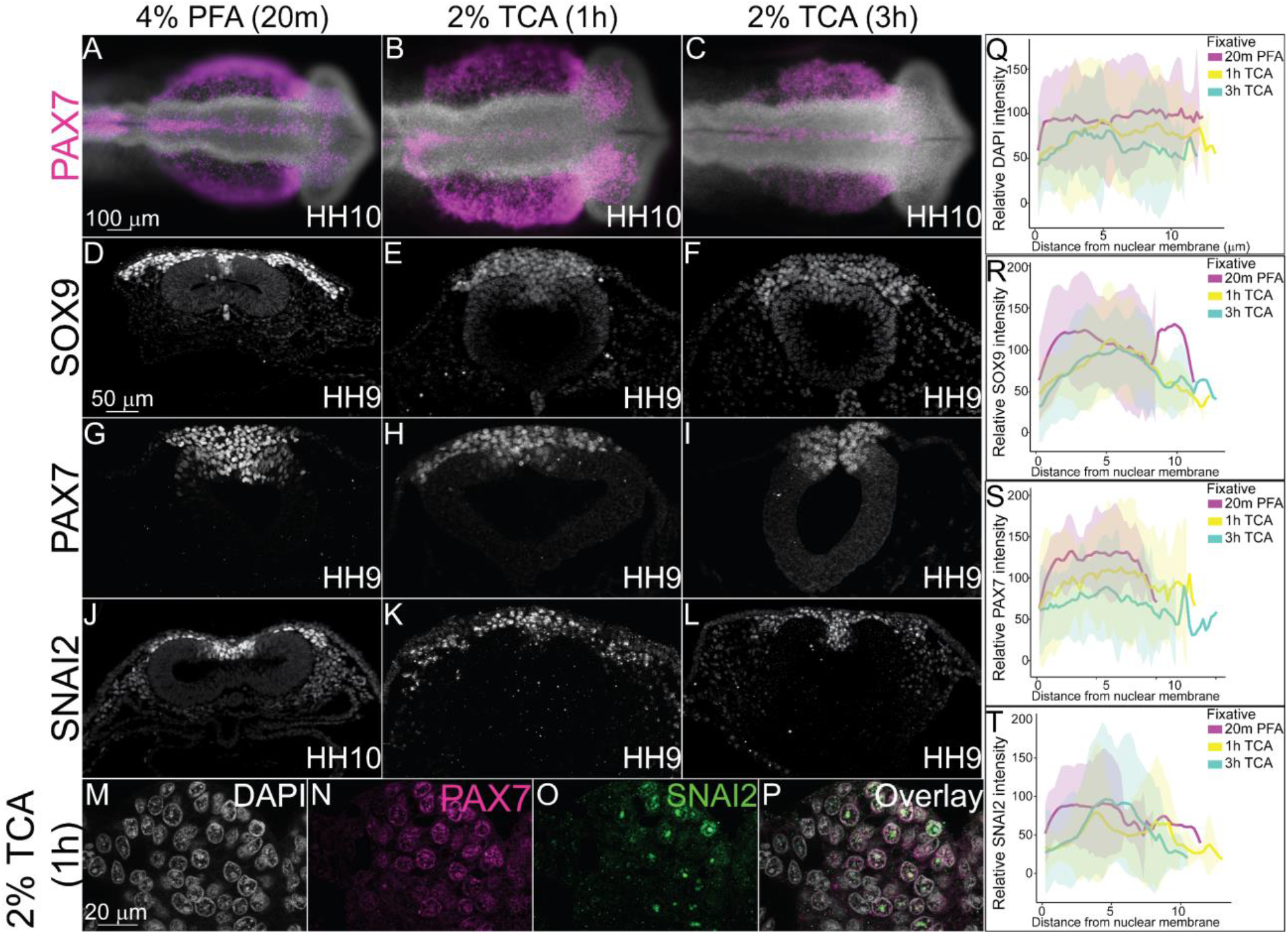
TCA and PFA fixation create differences in NC-specific transcription factor fluorescence levels across nuclei. IHC of definitive NC cell markers SOX9, PAX7, and SNAI2 in stage HH9-10 chicken embryos fixed in (A, D, G, J) 4% PFA for 20m, (B, E, H, K,) 2% TCA for 1h, and (C, F, I, L) 2% TCA for 3h. (A-C) Wholemount chicken embryos after IHC for PAX7 (purple) and DAPI staining (cranial region, anterior to the right). Transverse sections from embryos after IHC for (D-F) SOX9 (G-I), PAX7, and (J-L) SNAI2. (M-P) High magnification images of 2% TCA 1 hr fixed embryo nuclei after IHC for PAX7 and SNAI2 with DAPI staining. (Q) Graph showing relative fluorescence intensity of DAPI across the nucleus comparing the three fixative conditions shows relatively uniform fluorescence across the nuclei, *n*=5. (R) Graph showing relative intensity of SOX9 across nuclei comparing the three fixative conditions shows some variation in signal across the nuclei with variable peaks in intensity in PFA fixation, *n*=5. (S) Graph showing relative fluorescence of PAX7 across the nucleus comparing the three fixative conditions shows similar trends in intensity across nuclei in all fixatives, *n*=5. (T) Graph showing relative fluorescence of SNAI2 across the nucleus comparing the three fixative conditions show distinct fluorescence peaks in TCA fixatives, *n*=5. Fixation with 2% TCA for 1h increased visualization of nuclear spatial arrangement. The scale bar for the wholemount is 100 μm, the transverse section is 50 μm, and the high magnification transverse section is 20 μm.

PFA is the primary mode of fixation in avian embryos prior to performing IHC and it works effectively with short fixation times (Table 1). To determine how TCA fixation works for antibodies targeted to antigens in the nucleus, we used previously characterized antibodies against transcription factors paired box protein 7 (PAX7), SRY-Box 9 (SOX9), and Snail Family Repressor 2 (SNAI2) (Monroy et al., 2022). At HH10, the TCA-fixed wholemount embryos appeared larger than those fixed in PFA, but there were no significant differences in neural tube size (Figure 2A-C). SOX9, PAX7, and SNAI2 fluorescence was robust and pan-nuclear in PFA (Figure 2D, G, J). However, although the TCA-fixed embryos appearance in wholemount did not appear different than those that were TCA fixed, in section, SOX9 and PAX7 expression appeared diffuse and the brightness was weaker (Figure 2 E, F, H, I). In contrast, SNAI2 localization became more punctate and was localized to 2-3 regions within each nucleus in TCA fixation compared to a more uniform fluorescence in PFA fixation (Figure 2, compare J to K and L). In higher magnification images of sections from TCA-fixed embryos, the DAPI stain overlaps with diffuse PAX7 protein signal, but SNAI2 protein appears limited within the nucleus in all NC cells in which it is expressed (Figure 2M-P). In chicken embryos, PFA appears to be a more effective fixation method prior to IHC for robust fluorescence using antibodies against the transcription factors that were tested.

### Different fixation methods alter the signal intensity of microtubule subunit proteins

To determine how fixation methods affect cytoplasmic and cytoskeletal protein localization, we tested the various fixation treatments in HH9 chicken embryos and performed IHC using antibodies against Tubulin Beta 3 Class III (TUBB3), Tubulin Beta 2A Class IIa (TUBB2A), and Tubulin Alpha 4a (TUBBA4A). Signal for all three proteins was visible in all three fixative treatments (Figure 3). We identified that TUBB3 protein signal appeared more robust in NC cells compared to the NT fluorescence signal in 2% TCA fixation compared to 20% PFA (Figure 3D-F). For TUBB2A, with PFA fixation, the protein appeared localized to the non-neural ectoderm (NNE) and cranial mesenchyme with weaker expression in the NC and NT. With TCA fixation, the TUBB2A fluorescence in the non-neural ectoderm and cranial mesenchyme increased compared to the signal in the NC cells and NT to the point that signal is almost imperceptible in the NT (Figure 3G-I). After PFA fixation, TUBA4A appears to solely localize to the non-neural ectoderm, but after TCA fixation, the protein is visible in the cranial mesenchyme, NC cells and weakly in the NT (Figure 3J-L).

**Figure 3.**
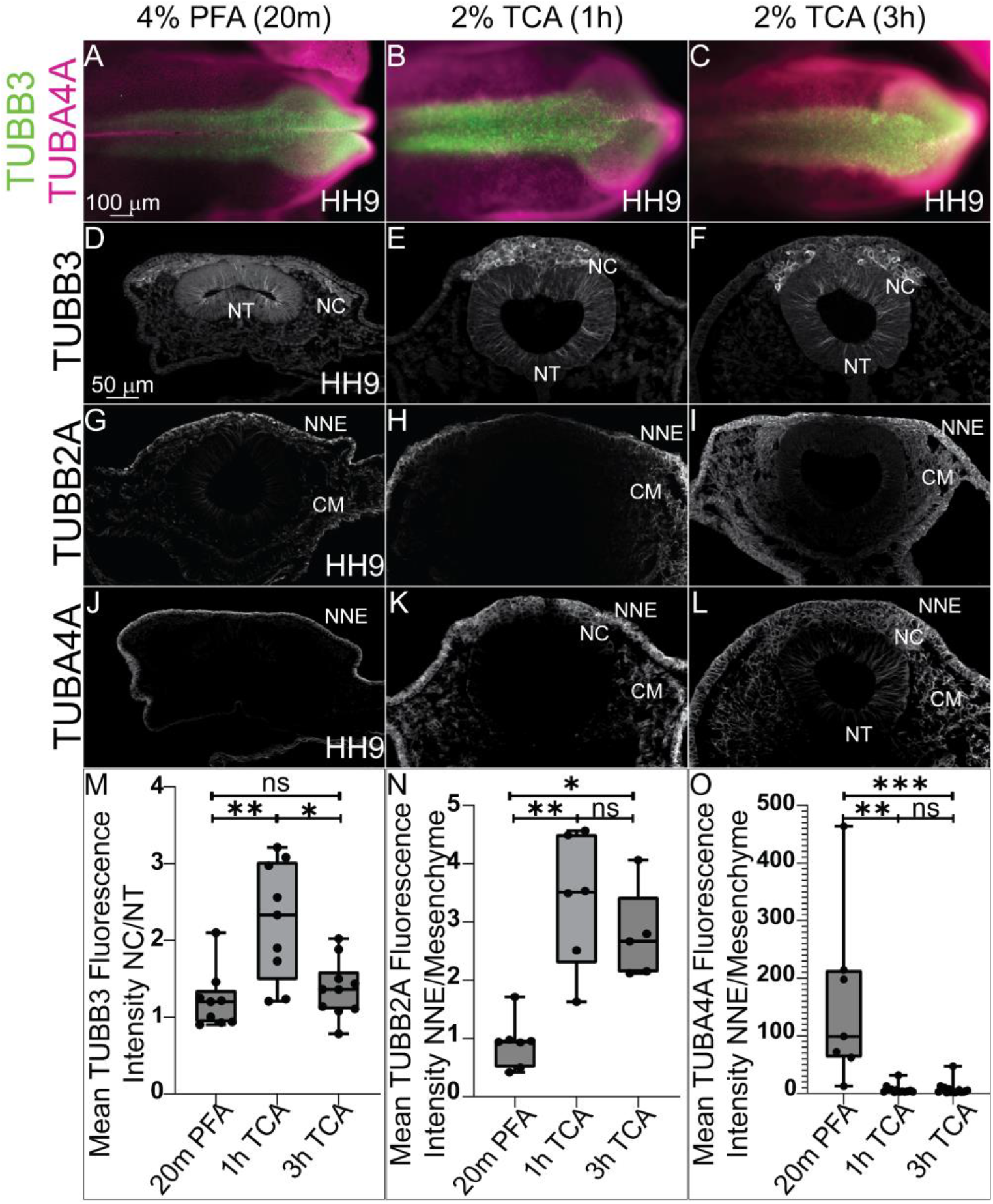
Differences in tissue-specific fluorescence levels of cytoskeletal proteins after TCA fixation. IHC using antibodies against the tubulin isotypes (A, D-F) TUBB3, (G-I) TUBB2A, and (A, J-L) TUBA4A at HH9 in (A-C) wholemount embryos and (D-L) transverse sections. The embryos were fixed in (A, D, G, and J) 4% PFA, in (B, E, H, and K) 2% TCA for 1h, and in (C, F, I, and L) 2% TCA for 3h. In whole embryo, across the three conditions TUBB3 is visible in the cranial dorsal side of the chicken embryo, and TUBA4A across the ectoderm. (D-F) TUBB3 signal appeared higher in NC cells after TCA fixation. (G-I) IHC for TUBB2A appeared to have a brighter mesenchymal signal after TCA fixation. (J-L) IHC for TUBA4A showed signal increase after TCA fixation. (M) The graph shows a ratio of mean fluorescence intensity of TUBB3 in NC cells compared to cells within the NT. The 1h TCA shows a significant increase in the NC cell fluorescence over cells within the NT compared to the 20m PFA and 3 hr TCA, (P ≤ 0.01 and P ≤ 0.05, respectively, n=9). (N) The graph shows a significant increase in TUBB2A fluorescence in the NNE compared to the cranial mesenchyme in 1 and 3 hr TCA compared to PFA (P ≤ 0.01 and P ≤ 0.05, respectively, n=7). (O) The graph shows a significant decrease in the NNE compared to the cranial mesenchyme in 1 hr and 3 hr TCA-fixed embryos compared to PFA (P ≤ 0.01 and P ≤ 0.001, respectively, n=7). NC= neural crest, NT= neural tube, NNE= non-neural ectoderm, CM= cranial mesenchyme. One-way ANOVA with the Kruskal-Wallis test was used to determine the significance differences in fluorescence intensity. The scale bar is marked in the first whole embryo and transverse section of the figure.

Quantifying changes in tissue-specific fluorescence intensity showed that TUBB3 fluorescence was significantly higher in NC cells compared to NT cells after 1 hr TCA fixation than it was after PFA fixation (Figure 3M) (P ≤ 0.01, n= 9) or 3 hr TCA fixation (P ≤ 0.05, n= 9). In addition, TCA fixation significantly enhanced TUBB2A intensity in the cranial mesenchyme compared to the NNE in both 1 hr and 3 hr treatments compared to PFA fixation (Figure 3N) (P ≤ 0.01 and P ≤ 0.05, respectively, n=7). The signal for TUBA4A appeared to be localized almost exclusively to the NNE after PFA fixation, but 1 hr and 3 hr TCA fixation increased the cranial mesenchyme and NT fluorescence signal and the intensity difference was reduced (P ≤ 0.01 and P ≤ 0.001, respectively, n=7). These data show that types of fixation are capable of altering the apparent signal in specific tissues.

### Different fixation methods affect cadherin protein tissue-specific intensity

The localization of N-cadherin (NCAD) and E-cadherin (ECAD) have previously been characterized in chicken embryos using PFA-fixation (Dady et al., 2012; Rogers et al., 2018). To determine if TCA fixation is also an efficient method prior to IHC to visualize these proteins, we tested the various fixation treatments in HH9 chicken embryos with antibodies against the two type-I cadherins. In PFA fixation, both cadherin proteins are expressed in the NT at HH9, but while ECAD is also expressed in delaminating NC cells and non-neural ectoderm, NCAD is absent from these tissues and instead present in the cranial mesenchyme (Figure 4A, D)(Dady et al., 2012; Rogers et al., 2018). In 2% TCA at both 1 hr and 3 hr fixations, ECAD localization remains in the same tissues (Figure 4B, C). We measured the relative fluorescence intensity in the developing non-neural ectoderm compared to the NT to determine if TCA fixation alters the signal intensity as it does in microtubule proteins, and although there was no difference in proportional intensity between PFA and the 1 hr TCA fixation (Figure 4G)(1.6 fold compared to 1.8 fold relative intensity difference, p=.06), in the 3 hr TCA fixation treatment, the non-neural ectoderm fluorescence was significantly higher than that in the NT (1.6 fold compared to 2.0 fold relative intensity difference, P ≤ 0.05, n= 6 TCA and n= 5 PFA).

**Figure 4.**
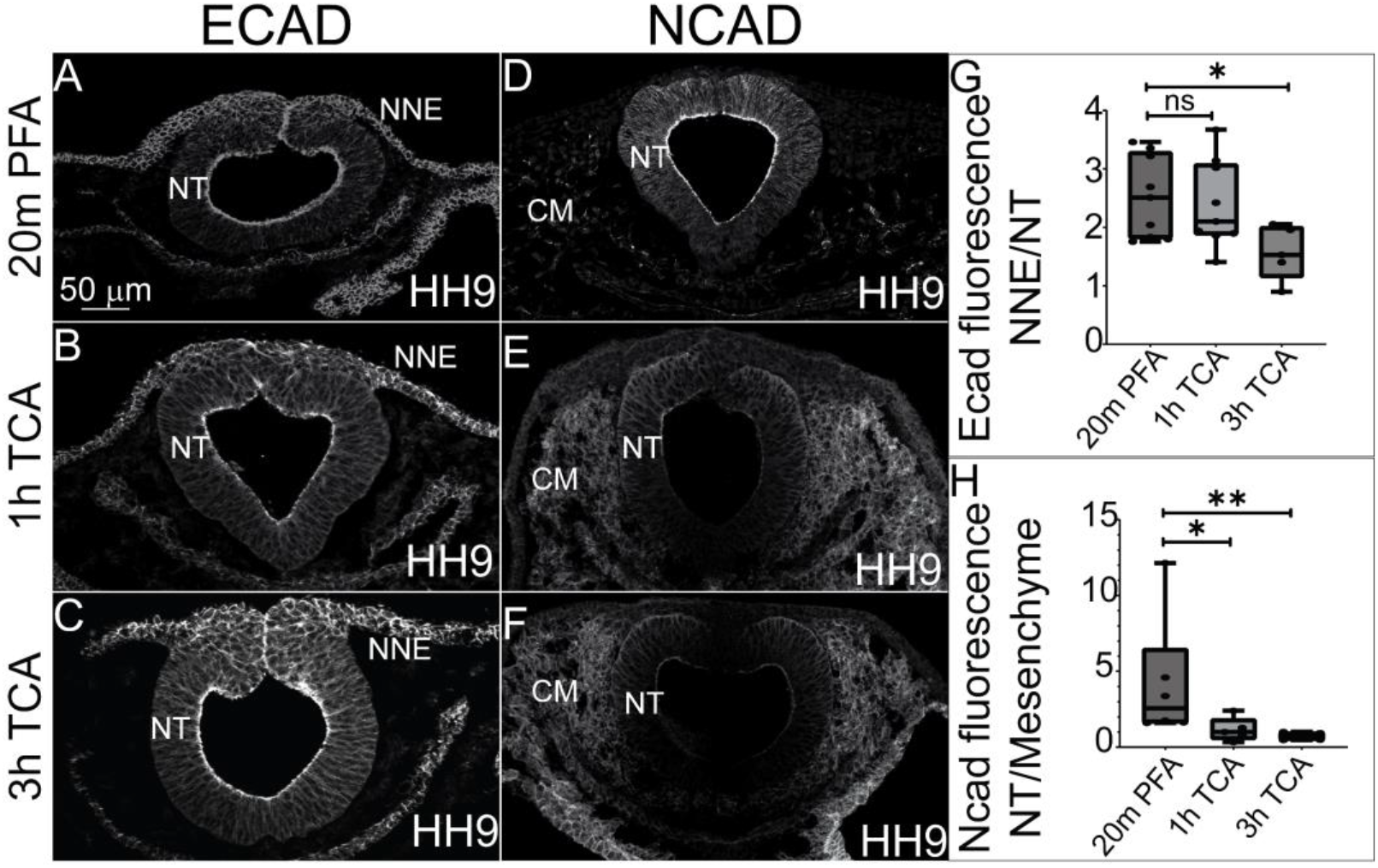
TCA and PFA fixation result in tissue-specific differences in fluorescence intensity of cadherin proteins. (A-F) Transverse sections comparing three fixative conditions prior to IHC for ECAD and NCAD in chicken embryos at stage HH9. (A-C) IHC for ECAD of embryos fixed in (A) 2% PFA, (B) 1 hr TCA, and (C) 3 hr TCA shows increased overall signal intensity after TCA fixation. (D-F) IHC for NCAD of embryos fixed in fixed in (D) 2% PFA, (E) 1 hr TCA, and (F) 3 hr TCA shows increased cranial mesenchyme and reduced neural tube signal intensity after TCA fixation. (G) The graph shows a significant decrease in the difference in intensity between the NNE and NT when comparing PFA to 3 hr TCA fixation (P ≤ 0.05, n= 6 TCA and n= 5 PFA). (H) The graph shows a significant decrease in the difference in fluorescence intensity in the NT compared to the cranial mesenchyme in embryos fixed for 1h in TCA (P ≤ 0.05, n=5) and 3h TCA (P ≤ 0.01, n=6) when compared to the PFA fixation. NT= neural tube, NNE= non-neural ectoderm, CM= cranial mesenchyme. One-way ANOVA with the Mann-Whitney test was used for analysis. The scale bar for transverse sections is 50µm.

In contrast to the relative lack of change in ECAD localization and minor changes in tissue-specific intensity with TCA fixation, the NCAD signal is strongly localized to the NT with weaker expression in the cranial mesenchyme using 4% PFA. In contrast, in 2% TCA fixation, the relative fluorescence intensity of NCAD significantly increases in the cranial mesenchyme after 1 hr (P ≤ 0.05, n=5) and 3 hr (P ≤ 0.01, n=5).

## 4. DISCUSSION

Despite their widespread use, studies have shown that over 50% of antibodies fail in one or more applications (Ayoubi et al., 2023). Thus, it is vital to validate that antibodies work properly before trusting them for characterization studies or functional applications. When using a new commercial antibody, researchers will often test various concentrations of the antibody, but may not validate the fixation method used to process the tissue beforehand. Here, we compared the effectiveness of PFA fixation to that of TCA fixation prior to IHC in whole chicken embryos using multiple previously validated antibodies. We identified that the type of fixation applied affected the resulting tissue morphology and the tissue-specific, and in some cases, subcellular visualization of proteins. The morphological changes are likely due to the different mechanisms by which PFA and TCA fix tissues. PFA covalently cross-links molecules, stabilizing tertiary and quaternary structures of proteins and hardening the cell surface (Kim et al., 2017). We observed that in PFA-fixed chicken embryos, tissue appeared more tightly packed with denser and less circular nuclei (Figure 1). In contrast, we found that TCA fixation resulted in larger and more circular nuclei (Figure 1). Rather than cross-linking proteins, TCA precipitates proteins by disrupting their encircling hydration sphere (Koontz, 2014). Unlike PFA, which maintains tertiary and quaternary structure, TCA denatures proteins to the point where their secondary structure is lost (Koontz, 2014). The cell shape changes we observed may be due to this precipitation of proteins within a cell, filling up space and rounding out the nuclear and cellular membranes.

These differences make each fixative type more ideal depending on the target epitope. Since TCA precipitates and denatures proteins, it makes hidden epitopes more accessible. In contrast, PFA is ideal for targeting structural epitopes as it maintains tertiary and quaternary structures. Here, we sought to understand how these various fixative methods affect immunohistochemical staining using antibodies for markers in multiple tissue types in chicken embryos. In contrast to PFA, which works well with short fixation times, such as the 15-20 minutes used for this study, TCA results in low signal at an equivalent fixation duration (data not shown). Thus, we employed one and three hours of TCA fixation, which generally led to similar outcomes of cellular morphology and signal intensity when compared to each other. By using chicken embryos as our model, we were able to use commercially available antibodies we have previously validated (Table 2). We identified that both PFA and TCA fixation allowed us to visualize proteins in their known locations, but we saw that some improved signal while others had detrimental effects.

We saw a marked difference in how TCA and PFA affected the visualization of nuclear markers. Nuclear markers had weaker fluorescence signal and appeared more compartmentalized within the nucleus in TCA-fixed embryos compared to PFA-fixed embryos (Figure 2). This result may be caused by actual subnuclear protein localization, or it may be due to the TCA precipitation of the target proteins within the nuclear compartment (Lakatos and Jobst, 1992; Rajalingam et al., 2009). In measuring fluorescence intensity of nuclear markers across nuclear membranes, we saw an increase in isolated peaks within nuclei, representing this compartmentalization (Figure 2). If this compartmentalization is biologically accurate, it is a method that could be used to visualize condensates within nuclei. Additionally, TCA fixation may allow us to compare the localization of multiple nuclear markers at once to see if their subnuclear localization differ at different phases of the cell cycle, for example. However, nuclear markers used following TCA fixation also tended to have a weaker signal compared to background, possibly due to precipitation.

PFA and TCA fixation also caused noticeable differences in the fluorescence intensity within specific tissues when used before IHC with microtubule subunits. We calculated this signal difference by measuring the ratio of fluorescence intensity between the most differentially fluorescent tissue types for a given marker between TCA and PFA. Past work showed TUBB3 is expressed in the neural tube at HH8 in chicken embryos, with a stronger level of signal at the dorsal side where the NC cells are present (Chacon and Rogers, 2019). Here, we see similar localization in HH9 chicken embryos, but one hour TCA fixation was optimal for showcasing the increase in the NC TUBB3 signal compared to the NT signal (Figure 3). Interestingly, TUBB2A and TUBA4A both displayed a marked difference in the ratio of NNE to cranial mesenchyme signal in the TCA fixation versus PFA fixation treatments, but in opposite directions. For TUBB2A, the NNE signal increased in TCA-fixed embryos compared to PFA-fixed embryos, increasing this ratio (Figure 3). Meanwhile, for TUBA4A, the cranial mesenchyme signal increased in TCA-fixed embryos compared to PFA-fixed embryos, decreasing this ratio (Figure 3). Thus, it is critical to test multiple fixatives for markers of interest even across similar protein types, as they may enhance signal in different tissue regions. Microtubule proteins are well known for their post-translational modifications which directly affects microtubule stability (Bar et al., 2022), and it is possible that these differently modified proteins are better targeted in one fixative versus the other.

We saw similar differences in cadherin protein IHC between TCA and PFA fixatives. In stage HH9 chickens, equivalent to our samples, ECAD localized to the NNE, NT, migratory NC cells, and developing gut (Rogers et al., 2018). While this localization remained consistent in our samples across various fixatives, we saw that the fluorescence intensity of the nonneural ectoderm increased compared to neural tube signal in tissues fixed with TCA three hours compared to the other fixative conditions (Figure 4). Similarly, NCAD displayed the expected localization for HH9 chickens to the neural tube, cranial mesenchyme, notochord, developing gut, and absence from the dorsal neural tube regardless of fixative type (Rogers et al., 2018). However, fluorescence intensity of NCAD in the neural tube was far higher than that in the cranial mesenchyme for PFA-fixed embryos compared to TCA-fixed embryos (Figure 4). This suggests that the type of fixative applied can affect the primary tissue in which a protein appears to be localized, and that issue may have far-reaching effects for individuals studying cell and developmental biology as those fields strongly rely on knowing spatiotemporal protein localization prior to studying protein function. Our results have implications for characterizing new antibodies that do not have published and validated expression models.

Future studies using IHC with traditional antibodies would benefit from fixation validation in addition to traditional antibody specification validations (knockdown, overexpression, Western blot, etc.) as some fixatives may improve visualization of proteins of interest. Comparing the 3 hr versus 1 hr TCA fixes to each other revealed that the one hour fixation is generally sufficient to alter tissue morphology and to reveal additional protein localization in the tissue samples. However, fixed tissues are not living tissues and as technologies become available, it would be important to visualize these cellular events *in vivo*. It may also be beneficial to compare additional fixation techniques such as alcohol-based fixation or antigen retrieval to see if these methods replicate or improve upon the outcomes from PFA or TCA fixation. As displayed in this paper, the method of fixation can affect how protein localization is visualized. While PFA revealed epitopes in most tissues, TCA-mediated protein denaturation may provide access to hidden epitopes in regions of the protein of interest that are inaccessible due to PFA cross-linking (Klockenbusch and Kast, 2010; Nadeau and Carlson, 2007). Here, we only tested these techniques in a single organism. However, the type of fixative used has been found to affect cellular and tissue morphology in other systems and animals including cell culture, goats, rats, and mice (Cox et al., 2006; Hirashima and Adachi, 2015; Paavilainen et al., 2010; Rahman et al.; Rezoana et al., 2022; Tu et al., 2011; Wang et al., 2016). The fixative type used should be optimized depending on the model system, type of protein, and expected localization. Though we focus on a single organism, we demonstrate that fixatives affect the visualization of numerous proteins in several cellular compartments. Our results demonstrate that methods can, and should, be tested for improved biological analyses and accurate demonstration of results in wholemount or in section.

## 5. AUTHOR CONTRIBUTIONS

Conceptualization: CDR, CVE, and TAL. Data Curation (experiments and imaging): CVE and TAL. Formal Analysis (cell counts, statistics, bioinformatics): CVE and TAL. Funding Acquisition: CDR, CVE, and TAL. Methodology: CDR, CVE, and TAL. Writing: CDR, CVE, and TAL. Supervision: CDR.

## 6. ACKNOWLEDGEMENTS

The authors would like to acknowledge the following funding sources: NSF CAREER award 2143217 and NIH R03DE032047-01 to CDR. TAL and CVE are funded in part by NSF GRFP awards. We would like to thank the members of the Rogers Lab at UC Davis and the UC Davis community for their discussions and input on this project.

## Notes

### Competing Interest Statement

The authors have declared no competing interest.

